# Disulfide chaperone knock-outs enable *in-vivo* double spin-labeling of an outer-membrane transporter

**DOI:** 10.1101/711663

**Authors:** T. D. Nilaweera, D. A. Nyenhuis, R. K. Nakamoto, D. S. Cafiso

**Affiliations:** University of Virginia

## Abstract

Recent advances in the application of EPR spectroscopy have demonstrated that it is possible to obtain structural information on bacterial outer-membrane proteins in intact cells from extracellularly labeled cysteines. However, in the *Escherichia coli* outer-membrane vitamin B_12_ transport protein, BtuB, the double labeling of many cysteine pairs is not possible in a wild-type K12-derived *E. coli* strain. It has also not yet been possible to selectively label single or paired cysteines that face the periplasmic space. Here we demonstrate that the inability to produce reactive cysteine residues in pairs is a result of the disulfide bond formation system, which functions to oxidize pairs of free-cysteine residues. Mutant strains that are *dsbA* or *dsbB* null facilitate labeling pairs of cysteines. Moreover, we demonstrate that the double labeling of sites on the periplasmic facing surface of BtuB is possible using a *dsbA* null strain. BtuB is found to exhibit different structures and structural changes in the cell than it does in isolated outer membranes or reconstituted systems, and the ability to label and perform EPR in cells is expected to be applicable to a range of other bacterial outer-membrane proteins.

**Statement of Significance:** EPR spectroscopy is an important method to characterize the structure and dynamics of membrane proteins, and recent efforts demonstrate that pulse EPR can be used to examine the extracellular surface of outer membrane proteins in live bacteria. In the present work, we show that pairs of cysteine residues in the Escherichia coli vitamin B_12_ transporter, BtuB, cannot be spin-labeled in wild-type strains, but can be labeled with the use of certain null mutants in the periplasmic disulfide bond formation, Dsb, system. These mutants also facilitate efficient spin-labeling of cysteines located on the periplasmic surface of BtuB. Distance measurements using pulse EPR provide evidence that the behavior of BtuB is different in the bacterial cell than it is in purified systems.

---

Gram negative bacteria are protected from the external environment by both an outer membrane (OM) and an inner membrane (IM), which in turn are separated by a periplasmic space. The OM of Gram-negative bacteria is populated by a dense network of lipopolysaccharide (LPS) molecules and outer membrane proteins (OMPs). These OMPs serve a variety of cellular roles, providing passive or active transport, adhesion, enzymatic activity, motility and other critical functions. TonB dependent transporters (TBDTs) are active transport proteins that are responsible for the uptake of a range of nutrients across the OM, including forms of iron, vitamin B_12_ and carbohydrates (1). These proteins are formed of a 22-stranded β-barrel surrounding a globular core (or hatch) domain. The extracellular surface of the barrel has long loops that have been implicated in transport, while the periplasmic face of the barrel is connected by short turns. The relative size of TBDT substrates is believed to necessitate large conformational shifts in the core domain. However, the mechanics of transport have not been elucidated, and the majority of the work on these proteins has been carried out on reconstituted systems, where transport has never been established.

The importance of the native environment to the organization and function of OMPs has recently become apparent. In vivo, OMPs are present at very high densities and may even be organized into distinct puncta on the cellular surface that are mediated by promiscuous interprotein interactions and hydrophobic mismatch (2–4). OMPs are also surrounded by LPS molecules, and in some cases directly interact with the LPS (5, 6). In the case of the *Escherichia coli* cobalamin transporter, BtuB, these LPS contacts have even been proposed to order the extracellular loops, alter the conformation of an exterior facing core loop, and shift the equilibria of an N-terminal energy coupling region termed the Ton-box (7). The Ton-box is located on the periplasmic face of BtuB, and in the reconstituted system it has been observed to unfold and extend into the periplasmic space in a substrate-dependent manner (8, 9). Recent efforts have demonstrated that both continuous wave (CW) and pulsed electron paramagnetic resonance (EPR) methods can be used to study BtuB in its native environment. Site-directed spin labeling (SDSL) followed by CW EPR was demonstrated for single-sites in intact cells and isolated outer membranes (10, 11). The ability to form doubly-labeled pairs between loops on the extracellular surface of BtuB for subsequent distance measurements through Double Electron-Electron Resonance (DEER) experiments has also been shown (10, 11).

Recently, we attempted to measure distances between core and barrel sites by pulse EPR on the extracellular surface of BtuB, but were not successful in obtaining cysteine specific labeling in cells using the RK5016 strain employed in the previous work. One explanation for this observation is the activity of the *E. coli* disulfide bond formation (Dsb) system. While the cytoplasm is highly reductive, precluding the presence of functional disulfides in cytoplasmic proteins, the periplasm is more oxidative and many periplasmic, IM, and OM proteins possess structural disulfide bonds. These bonds are formed during or after translation of the nascent protein through the SecYEG translocon via interactions with a catalytic Cys-X-X-Cys motif on the periplasmic protein DsbA (12, 13). Reduced DsbA is reoxidized by the IM protein DsbB, which is regenerated via the IM electron transport chain (12). For proteins with more than one disulfide, interconversion is made possible by the disulfide isomerase DsbC, which is in turn regenerated by another protein, DsbD (12). Modulation of the Dsb system is commonly employed for the production of eukaryotic proteins possessing multiple disulfide bonds in E. coli such as antibodies (14). In these cases, fusions of the target construct to DsbA or co-expression of DsbA with glutathione supplementation are typically employed to increase the rate of disulfide bond formation in the periplasmic space (15).

Distance measurements using pulse EPR typically require the introduction of two cysteine mutants into the protein of interest, which must remain reduced prior to spin labeling. In the present work, we demonstrate that single cysteines that can be labeled in wild-type cells become inaccessible to label when incorporated as cysteine pairs, thereby preventing labeling using a standard methanethiosulfonate spin label (MTSL) reagent. We also demonstrate that in these cases the use of strains lacking components of the Dsb system permit labeling of these cysteine pairs, and that both CW and pulse EPR are now possible. Moreover, spin labeling on the periplasmic surface of BtuB has not been possible in the wild-type strain, but mutants of the Dsb system permit labeling and pulse EPR measurements to be made on the periplasmic surface of BtuB. The ability to make these measurements enables a range of structural measurements on the native state and function of BtuB in cells, and it may also enable studies of other TBDTs and other proteins in the OM, such as the major trimeric porins OmpF and OmpC.

## Methods

### Cell lines and plasmids

pAG1 plasmid with wild type *btuB* gene and RK5016 *(-argH, -btuB, -metE*) were kindly provided by the late Professor R. Kadner, University of Virginia. *E. coli dsb* variant strains were obtained from the Coli Genetic Stock Center (Yale University, New Haven, CT). Strain RI89 possessed WT Dsb system (*araD139 Δ(araABC-leu)7679 galU galK Δ(lac)X74 rpsL thi phoR Δara714 leu*^+^), whereas strains RI90 (RI89 *dsbA::* Kan^r^), RI317 (RI89 *dsbB::*Kan^r^), and RI179 (RI89 *ΔdsbC::*Cam^r^) possessed null mutations in *dsbA, dsbB*, and *dsbC*, respectively.

### Preparation of whole cell samples

BtuB mutants (D6C-Q510C, S74C-T188C, V90C, T188C and V90C-T188C) were engineered using PIPE mutagenesis and verified by DNA sequencing (Genewiz, South Plainfield, NJ). The resulting *btuB* mutants and WT *btuB* containing plasmids were transformed into RK5016 or one of several *dsb* variant strains. In all cases, a single colony was used to inoculate Luria Bertani media and used to prepare glycerol stocks and stored at - 80° C. Precultures of minimal media (MM) (100 mM phosphate buffer, 8 mM NH_4_SO_4_, 2 mM sodium citrate, 200 μg/mL ampicillin, 0.2% w/v glucose, 150 μM thiamine, 3 mM MgSO_4_, 300 μM CaCl_2_, 0.01% w/v methionine, and 0.01% w/v arginine) were directly inoculated using small aliquots of the glycerol stock and were grown overnight at 37° C. Precultures were then used to inoculate MM main-cultures, which were grown with a minimum of three rounds of cell doubling to early-log phase at an optical density (OD) at 600 nm of 0.3. Culture aliquots of 50 mL were collected by centrifugation at 3260 x g for 10 minutes at 4 °C, and the cell pellet was first washed by resuspending in 10 mL of wash buffer (100 mM HEPES, pH 7.0) followed by pelleting the cells. Harvested cells were resuspended again in wash buffer and freshly prepared sulfhydryl reactive MTSL ((1-oxy-2,2,5,5-tetramethylpyrrolinyl-3-methyl)methanethiosulfonate) was added for labeling. The base dose of spin-label was 7.5 nmol/mL of cell culture at OD_600_ 0.3 and deviations in actual dosage are indicated in the corresponding figures. Labeling proceeded in the dark for 30 minutes, after which the cells were resuspended in 100 mM HEPES, pH 7.0 for a 30-minute wash step. This was followed by an additional 30-minute wash in a D_2_O buffer of 100 mM MES, pD 5.5, and collection of the cells by centrifugation at 3260x g for 6 minutes. It should be noted that the growth times used in the current procedure are shorter than those used in previous procedures (10, 16). This shorter growth regime maintains robust protein expression, while maintaining a viable cell population where death and lysis are at a minimum (17).

### Preparation of samples on the periplasmic face of BtuB

Growth and spin labeling proceeded as above, with the exception that many of the later steps were shortened to reduce label reduction in the periplasmic environment. The spin-labeling step proceeded for 30 minutes and was followed by a single resuspension step into 100 mM HEPES, pH 7.0. For labeling of the D6C-Q510C sites, all buffers were also supplemented with 2.5% glucose, which was found to reduce reduction of the MTSL label.

### Treatment of cells with DTT

After growth to an OD_600_ value of 0.3, RK5016 cells were resuspended in 100 mM HEPES, pH 7.0, with 100 mM dithiothreitol (DTT) in an attempt to facilitate labeling of the double cysteine mutants (Fig. S1). The cells were incubated for 10 minutes at 37 °C after which labeling and washing steps proceeded as before.

### Outer-membrane (OM) preparation

The desired single or double cysteine mutant BtuB expressing cells were grown as above in which the main cultures were grown for 8 hours instead of OD_600_ 0.3. Cells were pelleted at 6,080 x g for 10 minutes and the pellets were resuspended in 10-15 mL of 0.1 M HEPES, pH 8.0 (HEPES buffer), supplemented with 40 µM of 4-(2-aminoethyl) benzenesulfonyl floride hydrochloride (AEBSF) and 0.025 U of Benzonase Nuclease. Cells were disrupted using a French Press, and the cell debris pelleted by centrifugation with an SS-34 rotor at 17,210 x g for 20 minutes. The resulting supernatant was then spun at 118,730 x g for 60 minutes to pellet the membrane fractions, and the pellets were resuspended in HEPES buffer supplemented with 1% sarkosyl. The resuspension was then incubated at 37 °C for 30 minutes, followed by a second 60-minute spin at 118,730 x g. The pellet was resuspended in HEPES buffer to a final volume of 5 mL, and 0.75 to 1.5 μmol/mL MTSL was added for spin labeling, which proceeded for 2 hours in the dark. Following spin labeling, the sample was spun at 125,750 x g for 20 minutes. The supernatant was discarded, and the pellet surface washed 3 times with HEPES buffer. The pellet was resuspended and spun in a tabletop-ultracentrifuge at 156,424 x g for 20 minutes. The pellet was surface washed twice with HEPES buffer, and manually resuspended into the same buffer using a pipette tip. The tabletop spin-wash-resuspend cycle was repeated two or three additional times, after which the pellets were resuspended in minimal volume HEPES buffer and flash frozen in liquid nitrogen for storage.

### EPR spectroscopy

For continuous wave (CW) EPR spectra, a volume of 6 uL of cell pellet or isolated membranes was loaded into 0.84 × 0.6 mm quartz capillaries (VitroCom, Mountain Lakes, NJ). Pellets were spun down with a hand-crank centrifuge. EPR spectra were recorded at room temperature at X-band using a Bruker EMX spectrometer with an ER 4123D dielectric resonator using a sweep width of 100 Gauss, a modulation amplitude of 1 Gauss, and 2 mW incident microwave power. Data were collected as additive averages of n-scans and were normalized by their second integral where indicated.

For double electron-electron resonance (DEER), 16 μL of concentrated cell pellet or isolated membranes was combined with 4 μL of deuterated glycerol for the apo samples and loaded into quartz capillaries (1.5 ID × 1.8 OD 100 mm length). For samples containing substrate, an additional 2 μL of cobalamin was added to a final concentration of 100 μM. All DEER experiments were performed on a Bruker E580 spectrometer operating at Q-band with an EN5107D2 dielectric resonator, SpinJet AWG, and a 10 W solid-state amplifier. Experiments were run at 80 K using the standard dead-time free 4-pulse DEER experiment, with an 18 ns π/2 pulse and 36 ns π pulses (18). All pulses were rectangular. The separation between observe and pump frequencies was 75 MHz.

DEER data were analyzed using LongDistances version 888 (Christian Altenbach, UCLA). Data were loaded into LongDistances and the phase and start time were determined automatically in the program. Five or six data points were truncated from the end of each trace to suppress 2+1 pulse artifacts, and the background was fit to a variable dimension stretched-exponential. Distance distributions were extracted using the Model Free mode.

### Molecular structures

Protein visualizations were generated using Pymol (19). The schematic representation of the Dsb system was generated via Pymol and BioRender. The DEER and CW data were plotted using python and the plotly.py, matplotlib, and numpy software packages.

## Results

The structure of BtuB in the apo state (PDB ID: 1NQG) is shown in Figure 1 with site V90 in the core domain and site T188 in the 2nd extracellular loop that connects strands 3 and 4 in the barrel. In the crystal structures of apo and substrate-bound BtuB (PDB ID: 1NQH), the loop containing residue 90 undergoes a substrate-dependent conformational change, with a short helical segment becoming unstructured upon substrate binding (20). Moreover, molecular dynamics (MD) simulations of BtuB predict that this conformational shift should be dependent upon the native environment (7).

**Figure 1.**
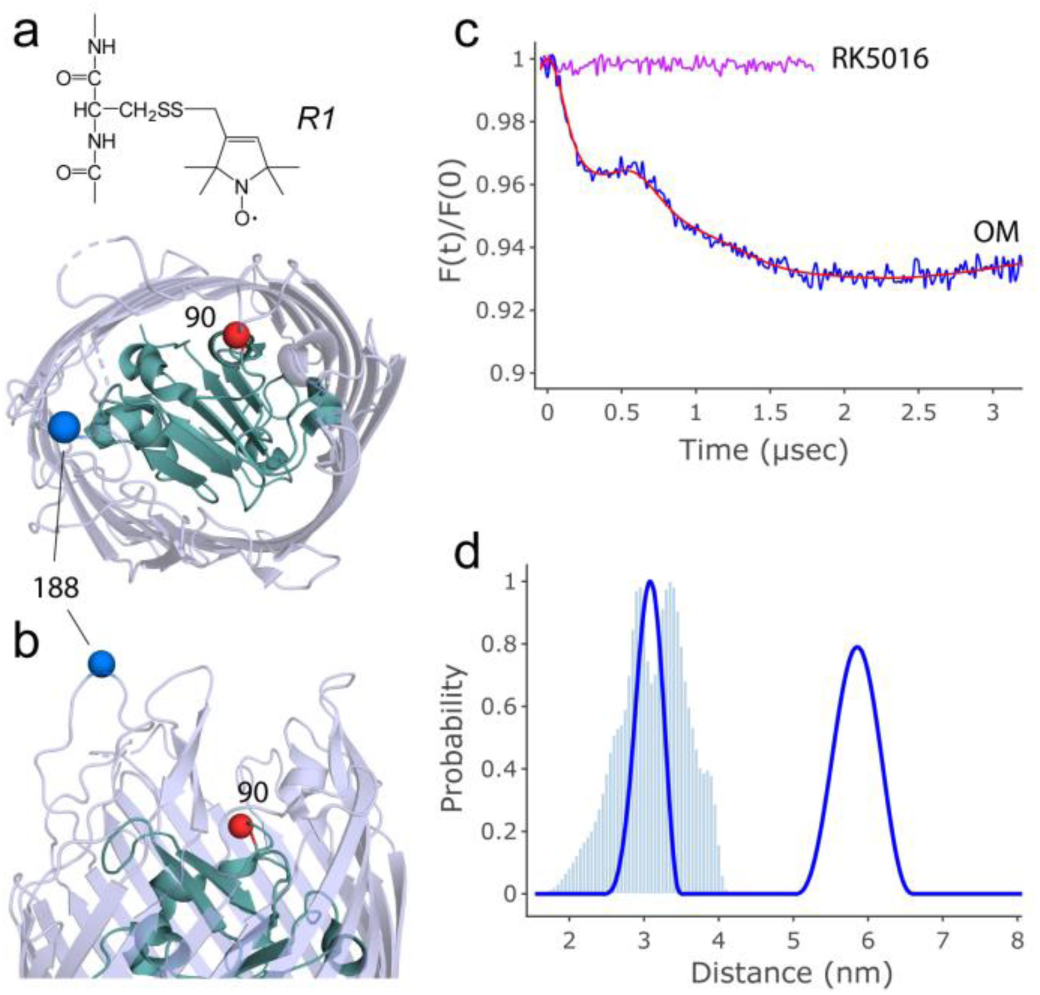
Preliminary DEER measurements at sites V90R1 and T188R1 on the extracellular surface of BtuB in cells and OM preparations. (**a**) The spin labeled side chain R1, which is a result of the reaction of the MTSL side chain with a reduced cysteine. (**b**) Top and side views of BtuB (PDB ID: 1NQG) showing the positions of residue V90 (pink) on an upper hatch domain loop and residue T188 (blue) on loop 2. (**c**) Dipolar traces obtained after background correction of DEER data obtained for BtuB 90-188 in isolated OMs (blue) and RK5016 whole cells (top trace, purple). The red trace represents the fit to the OM data. The absence of modulation in the whole cell sample suggested a lack of intramolecular interactions. (**d**) The distance distribution obtained for the isolated OM sample shows a short distance for the V90R1-T188R1 pair that agrees well with the predicted crystal structure (blue histogram shows the prediction from MMM). A longer distance component is also present that may represent intermolecular organization of BtuB on the cell surface.

To determine whether structural changes can be detected in this region, the doubly-labeled BtuB V90C-T188C pair was overexpressed in RK5016 cells, and DEER was run on an isolated OM preparation (Fig. 1c and d, blue). In the OM preparation, two peaks were observed for this pair, with a short-component at 3 nm and a longer distance between 5 and 6 nm. Although not well-defined, this longer distance may be the result of intermolecular interactions between neighboring BtuB molecules in the OM (see Discussion). The short distance aligns favorably with the V90R1-T188R1 intramolecular distance (Fig. 1d) predicted from the apo crystal structure using the program MMM (21).

Labeling was then attempted for the V90C-T188C pair in whole RK5016 cells. Surprisingly, the resulting dipolar trace showed no apparent modulation depth (Fig. 1c, top trace, purple), indicating a total absence of intramolecular interactions and a lack of specific protein labeling. Since both cysteine sites could be labeled individually in RK5016 cells (10), and since the protein was successfully doubly-labeled in the isolated OM, the state of the protein must be altered on the cell surface.

The redox potential of the periplasm is tightly controlled, and one explanation for the result in Fig. 1c was that oxidation had resulted in the formation of an intramolecular V90C-T188C disulfide bond. Attempts to vary the level of the MTSL reagent and introducing DTT (Fig. S1) generated either excess aqueous free label or a non-specific background signal but failed to facilitate labeling of this cysteine pair (see Supporting Information). In total, with variations in MTSL and DTT concentrations, we attempted to label the RK5016 cell line 13 times at sites V90C-T188C and 17 times in RK5016 at sites S74C-T188C. In only one case, for the 90-188 pair, were we able to obtain a small DEER signal with a modulation depth of 1.5% that yielded a short distance roughly consistent with the expected distance (Fig. S2).

### The use of dsbA or dsbB null mutant strains allows labeling of the V90C-T188C spin pair in cells

To test whether the Dsb system was preventing efficient labeling of BtuB in the cell by the MTSL reagent, null mutant strains were obtained for *dsbA, dsbB*, and *dsbC* alongside the corresponding WT strain, referred to here as *dsbA*^−^, *dsbB*^−^, *dsbC*^−^ and WT *dsb* respectively. The *dsbA*^−^ strain was selected for initial screening based on the direct interaction of DsbA with template cysteines. BtuB was then overexpressed in whole *dsbA*^−^ cells and labeled with varying concentrations of spin label (Fig. S3). In this cell line, low concentrations of spin label produced a lineshape consistent with specific labeling of sites V90R1 and T188R1, whereas higher concentrations resulted in significant aqueous label and an increase in non-specific labeling that may reflect nonspecific adsorption of the label to the cell surface. As a control, labeling was attempted on the *dsbA-* strain in the absence of BtuB V90C-188C expression. As seen in Fig. S4, no specific labeling is observed.

The level of MTSL reagent was chosen to maximize the apparent signal without introducing a significant free spin signal. This was then used for the remaining strains in order to verify that the signal observed was due to the loss of disulfide formation through the removal of DsbA. The resulting CW spectra are shown in Fig. 2. Both *dsbA*^−^ and *dsbB*^−^strains produced labeling of overexpressed BtuB V90R1-T188R1 (blue trace) that was significantly above the background observed for WT BtuB overexpressed in the same strains (red trace, Fig. 2a). In contrast, BtuB V90C-T188C overexpressed in *dsbC*^−^ or the WT *dsb* strains produced no apparent increase in labeling compared to WT BtuB. This result indicated that only removal of the DsbA protein, which is directly responsible for formation of the intramolecular disulfide bonds, or removal of DsbB, which is required for reoxidation of DsbA, permitted efficient labeling of the protein on the OM surface. It should be noted that each of the cell lines in Fig. 2 is expressing BtuB V90C-188C (see Fig. S5).

**Figure 2.**
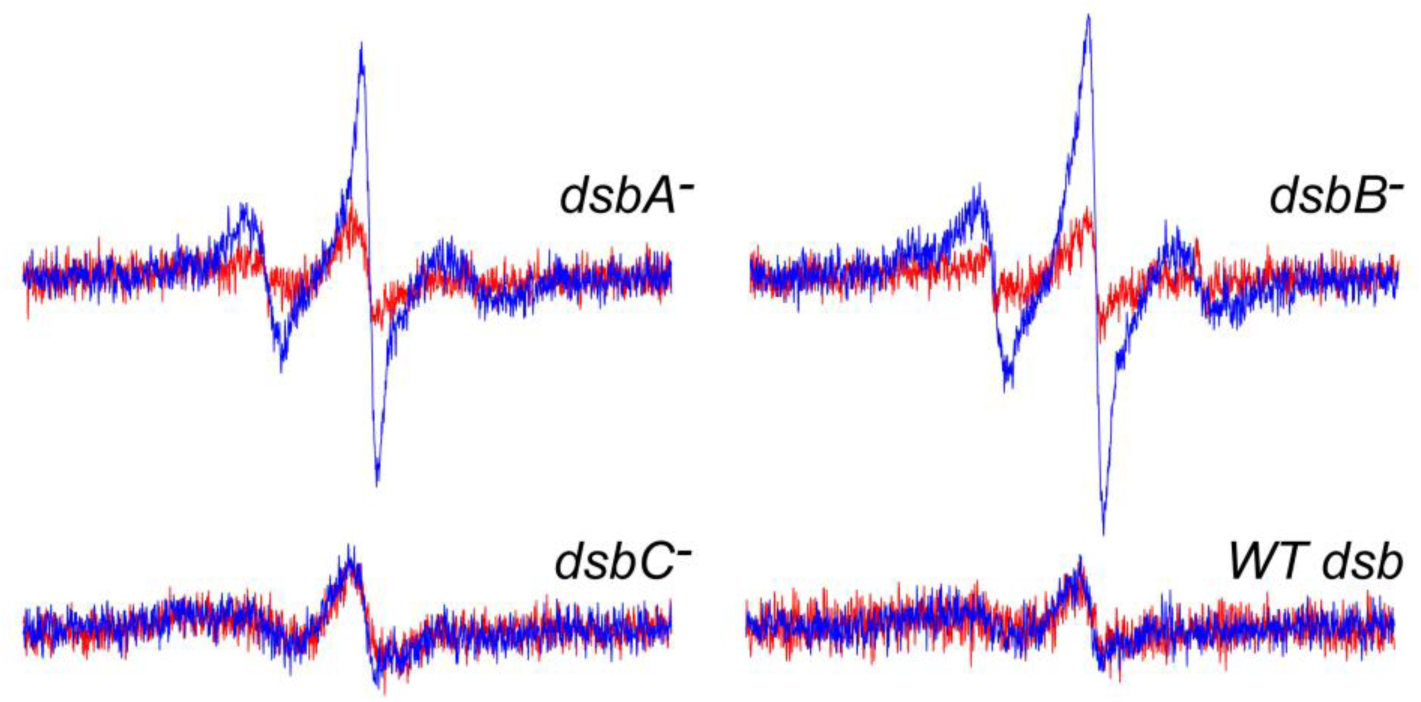
Results of attempting to label the BtuB V90C-T188C pair in *dsb* knock-out cell lines. CW spectra obtained for BtuB V90C-T188C in *dsbA*^−^, *dsbB*^−^, or *dsbC*^−^ cell lines, or in the WT *dsb* strain. The BtuB V90R1-T188R1 spectra (blue) are overlaid against spectra obtained from overexpressing WT BtuB in the same strain (red). Only *dsbA*^−^ and *dsbB*^−^ strains display labeling above the WT BtuB baseline. The X-band spectra were obtained from a single 21 second scan of 100 Gauss.

Pulse EPR data were collected for spin-labeled V90C-T188C using the *dsbA*^−^ and *dsbB*^−^ strains as well as the *dsbC*^−^ strain, which was not expected to produce a significant DEER signal. As shown in Fig. 3, both the *dsbA*^−^ and *dsbB*^−^ strains produced traces having a significant modulation depth, and the resulting distance distributions revealed distinct short components between 2 and 3 nm, as seen in the OM trace (Fig. 1). By contrast, the *dsbC*^−^ strain produced a much smaller dipolar trace having approximately 1 percent modulation depth, indicative of only a few percent of doubly-labeled BtuB molecules on the OM surface. Thus, the DEER data are consistent with the CW spectra and indicate that the ability to double-label BtuB V90R1-T188R1 is dependent on preventing DsbA/DsbB mediated disulfide bond formation.

**Figure 3.**
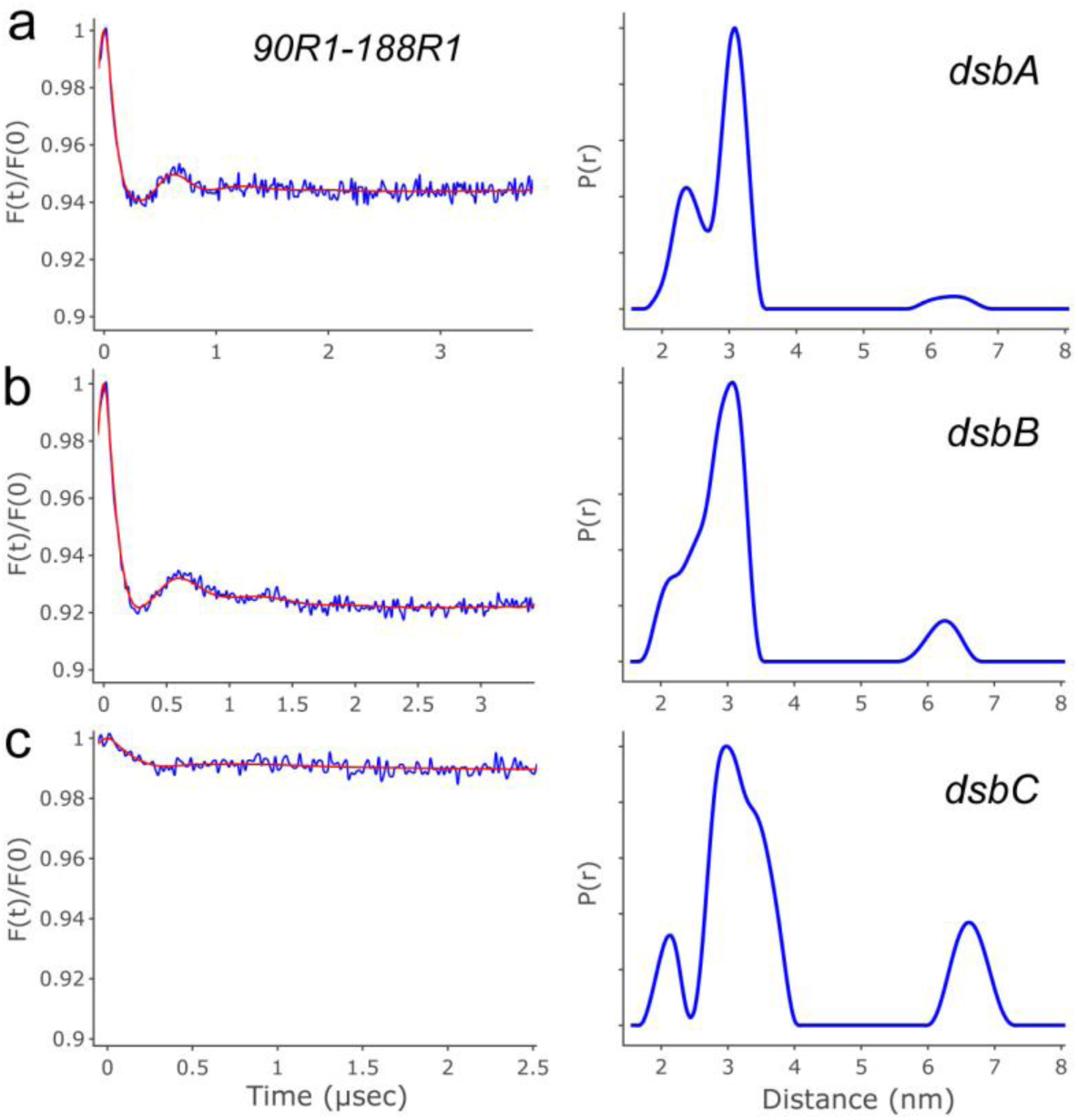
DEER data obtained for the BtuB V90C-T188C pair overexpressed in the three dsb system null mutant strains. The dipolar evolution and best fits (red) are shown on the left panels and distance distributions are shown on the right for (**a**) *dsbA*^−^, (**b**) *dsbB*^−^, and (**c**) *dsbC*^−^. As with the CW results, the *dsbA*^−^ and *dsbB*^−^ strains produce significant modulation depths and show short-distance components consistent with the predicted distance from the apo structure and with the previously shown results in OM isolates. The longer-distance peak is suppressed in these strains, which may indicate a decrease in protein expression level. By contrast, the *dsbC*^−^ strain has a markedly smaller depth consistent with only a small percentage of proteins having two accessible cysteines for spin-labeling. The resulting distance distribution was poor in quality, but still contained a short-distance component near the expected distance, indicating that the small interacting population was still BtuB V90R1-T188R1.

Compared to the single short-distance peak observed for BtuB V90R1-T188R1 in isolated OMs, the distribution obtained in *dsbA*^−^ cells showed a split-distance at a comparable smoothing factor, with peaks at 2.4 and 3.1 nm. To investigate this further, the experiment was repeated with 100 μM cobalamin (Fig. 4a). The populations of these two peaks invert with cobalamin addition, so that the distribution is dominated by the shorter 2.4 nm peak. A conformational change is also predicted at this site from the apo and substrate bound crystal structures (histograms, Fig. 4a), but the distance change predicted from the crystal structures is not as large as that observed using DEER. Surprisingly, this substrate-dependent change is not seen in the OM sample.

**Figure 4.**
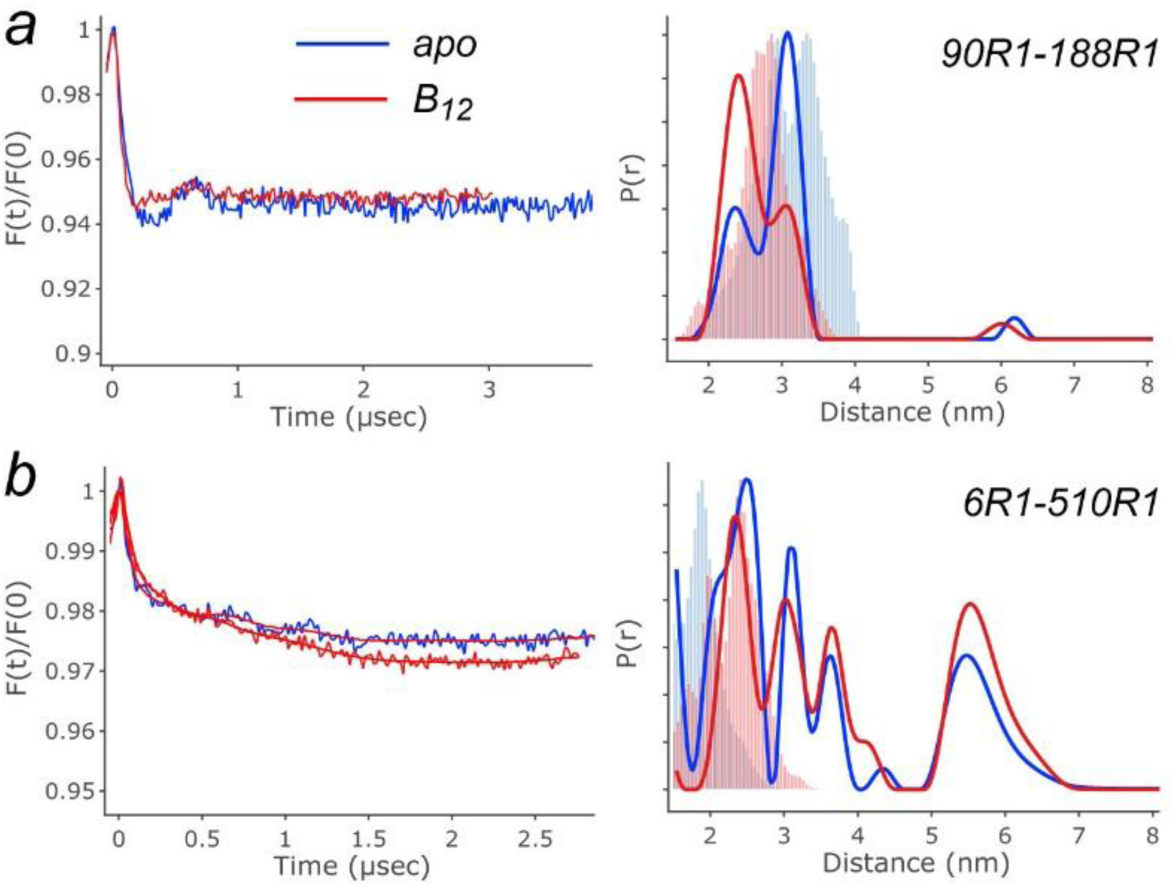
Comparison of DEER data in the apo state and after the addition of 100 μM cobalamin for BtuB V90R1-T188R1 on the extracellular surface, and BtuB D6R1-Q510R1 on the periplasmic face. (**a**) The addition of cobalamin to BtuB V90R1-T188R1 causes a shift in the observed peaks from a dominant mode at 3.2 nm in the apo condition to 2.4 nm with the addition of cobalamin. This may represent a disordering of the loop containing site V90 in response to substrate binding and is more apparent in the experimental data than in the crystal structure predicted distances for the apo (blue histogram) and cobalamin bound (red histogram) structures. (**b**) On the periplasmic surface, the D6R1-Q510R1 pair has a short distance component consistent with a folded Ton-box element observed in the apo (blue histogram) and cobalamin bound (red histogram) structures. There are also two intermediate distances that indicate partial unfolding or extension of the Ton-box, which are consistent with our previously observed results in reconstituted systems. No change is observed with substrate, indicating that the Ton-box may be constitutively unfolded in the native system. The long-distance component in this trace may represent an intermolecular interaction between neighboring proteins, or an additional longer mode of Ton-box extension.

In addition to V90C-T188C and 74C-T188C, pairs of cysteines at other sites within BtuB can only be labeled with the use of a Dsb mutant strain. Several examples of additional sites that require the *dsbA-* strain for labeling are shown in Fig. S6. We have not made an exhaustive study of where pairs of cysteines are cross-linked within BtuB by the Dsb system, but the high efficiency of this system may reduce the availability of cysteine pairs to the MTSL reagent at many sites.

### The dsbA^−^ strain permits labeling and pulse EPR on the periplasmic face of BtuB

In previous work, it was not possible to spin label cysteines on the periplasmic face of BtuB and obtain DEER data in cells, presumably due to the reductive potential of the periplasm (22). Since the Dsb system may have played a role in preventing labeling, we decided to re-examine labeling of the periplasmic surface using one of the *dsb* null mutant strains. Cysteines were incorporated into a core-barrel cysteine pair, D6C-Q510C, which was previously characterized in a membrane reconstituted system. This pair, between site D6 at the N-terminal end of the energy coupling Ton-box and Q510 in a periplasmic turn, previously yielded distance components in the reconstituted system consistent with a substrate-dependent unfolding and extension of the Ton-box into the periplasm (9).

When *dsbA*^−^ cells overexpressing BtuB D6C-Q510C were labeled with MTSL, a reduction of the spin label was still observed resulting in a rapid loss of the CW signal. Thus, the periplasm of the *dsbA*^−^ cells was still reducing. However, even with this reduction, it was possible to prepare cell samples that yielded DEER data. Shown in Fig. 4b are DEER data for the D6R1-Q510R1 spin pair in cells. The modulation depth of the dipolar trace is reduced compared to the extracellular V90R1-T188R1 pair, presumably due to label reduction that occurs in the periplasm. However, using this *dsb* null strain, DEER data are of sufficient quality to resolve several main peaks in the distance distribution. The first peak near 2.5 nm is consistent with the predicted distances from the apo (blue) and cobalamin-bound (red) crystal structures, where the Ton-box is folded into the hatch domain. The next two peaks, near 3 and between 3 and 4 nm, indicate that there is a partial periplasmic extension of the Ton-box. These two distances were observed previously in the reconstituted system upon the addition of substrate (9). Finally, the peak between 5 and 6 nm may be due to intermolecular interactions of BtuB or it may represent a further extension of the Ton-box element. Remarkably, in contrast to the reconstituted system, no extension or change in the Ton-box is observed after cobalamin addition, and the Ton-box appears to be permanently unfolded in the cell.

## Discussion

The formation of disulfide bonds in Gram-negative bacteria is tightly controlled by the Dsb system, and an active Dsb system is advantageous for cellular survival, as many periplasmic proteins are reliant on functional disulfide bonds (23). The system can be exploited in the production of complex eukaryotic target proteins with multiple disulfides (15), and it has sufficient potential to oxidize pairs of cysteines in overexpressed proteins even when one of the two *dsbA* promoters is removed (24). However, when a pair of reduced cysteine residues is required for spin-labeling on the cell surface, as is the case with many pulsed-EPR DEER experiments, a robust Dsb system may not be desirable.

For the labeled sites examined here, cysteine mutants in the hatch and barrel of BtuB could be singly-labeled in whole cells but could only be double-labeled in an isolated OM preparation (Fig. 1). In whole cells, there was a lack of specific labeling and a complete absence of intramolecular dipolar coupling in the DEER signal. Since dipolar coupling could be observed in the DEER signal from an isolated OM preparation derived from these cells, the protein was being inserted into the OM. This suggested that the state of BtuB had been modified in the cell, but that the OM isolation and labeling protocol rendered the protein amenable to labeling. There may be several explanations for our ability to label BtuB in the OM preparation but not the cell preparation. First, labeling of the OM preparation typically proceeded for several hours with excess MTSL reagent. If the cysteine mutants were linked in a disulfide bond, this protocol might have allowed the MTSL to act as its own reducing agent prior to labeling. Another explanation is that cell-disruption during the OM isolation procedure might have released reductive potential from the cytosol that reduced the disulfide, thereby permitting labeling by the MTSL.

To test the idea that the Dsb system was acting to cross-link pairs of cysteines in the whole cell (see Figure 5), we utilized several null mutant strains lacking proteins in the Dsb system. As seen in Figure 2, only DsbA and DsbB, which are directly involved in formation of template disulfide bonds, successfully produced CW EPR spectra above background labeling levels and yielded significant modulation depths in the DEER data. As expected for a target with only two cysteine mutants, the *dsbC*^−^ strain produced a much smaller modulation depth in the DEER data and failed to produce a significant deviation from the WT baseline in the CW EPR spectra.

**Figure 5.**
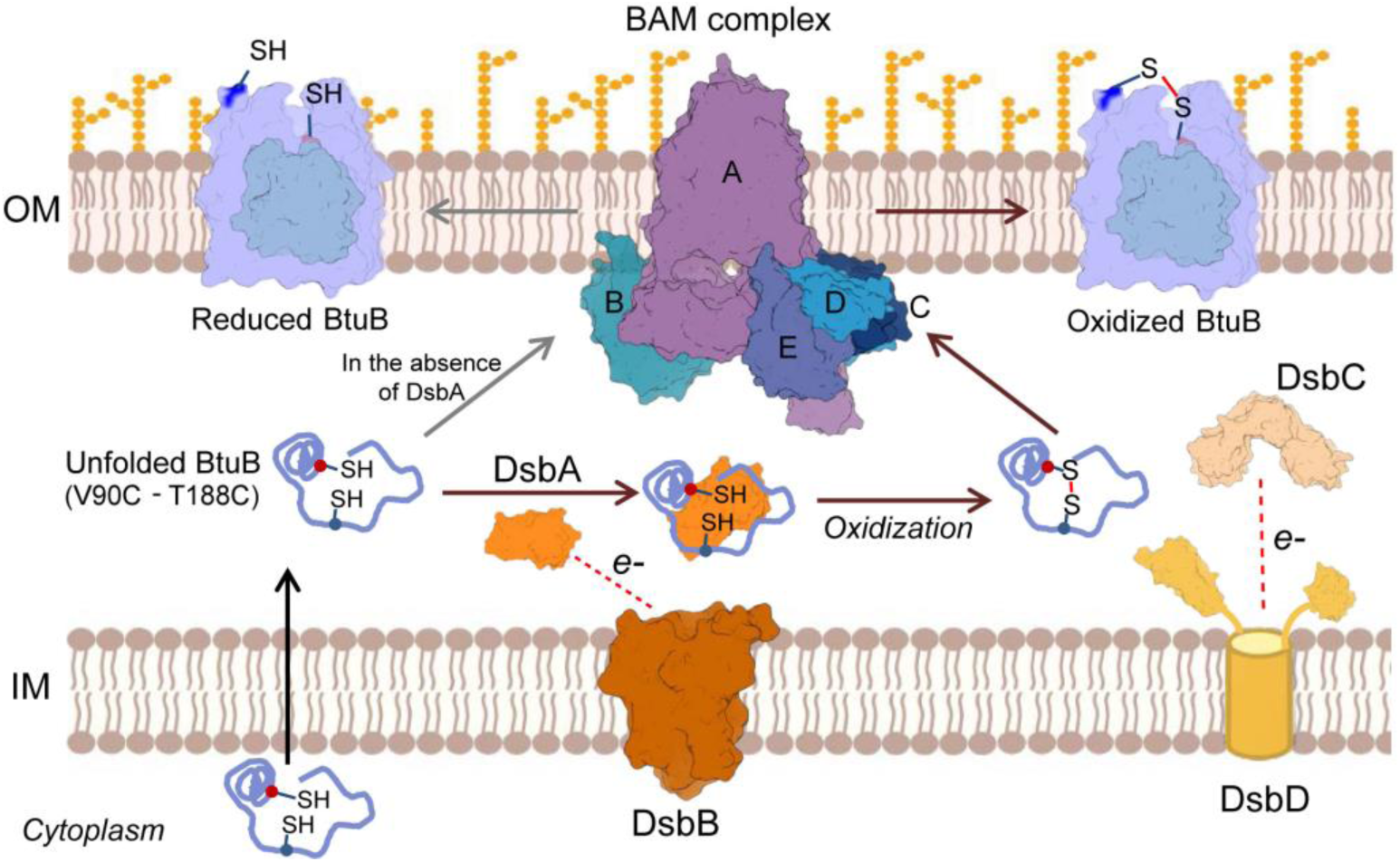
The Dsb system functions to cross-link pairs of cysteine residues in proteins such as BtuB that are destined for the outer-membrane (23). Electrons are extracted from BtuB by DsbA (PDB ID: 1A2L), which delivers them to DsbB (PDB ID: 2HI7), ultimately sending them into the electron transport chain. DsbC (PDB ID: 1EEJ), which had a minimal effect on the state of the V90C-T188C cysteine pair in BtuB (Figs. 2 and 3c), acts as an isomerase. DsbC is recharged by DsbD ((PDB ID: 1L6P and 2FWE) in the IM. BtuB (PDB ID: 1NQG) is folded and inserted into the OM at the BAM complex (PDB ID: 5D0O).

In previously published work, it was possible to acquire DEER data in intact *E. coli* between two sites in the extracellular loops; in particular, between residue T188R1 in loop 2 and residue G399R1 in loop 7 (10). Unlike the results presented here, this was possible without the use of *dsbA*^−^ or *dsbB*^−^ strains. There may be several reasons why labeling of the T188C-G399C pair was possible. First, the ability to label this pair may be due to the greater distance between these sites, both spatially and sequentially, which might have made a crosslink more difficult to form. Second, these loops are on opposite sides of the barrel, and formation of this crosslink prior to insertion might have blocked proper protein folding, leading to premature degradation of those proteins possessing the crosslink. If some percentage of nascent BtuB escaped the Dsb machinery, it would still be incorporated, albeit at a slower rate, and might have been labeled and detected by DEER. The longer growth time that was used prior to labeling in this earlier work may have helped facilitate this measurement.

A long-distance component appears in some of the data presented here that is consistent with an intermolecular (BtuB-BtuB) interaction. The appearance of this component is not unexpected and is consistent with the tight punctate organization of OM proteins observed in fluorescence studies of LamB, and recently demonstrated for BtuB and Cir bound to fluorescently tagged colicins (3, 4, 25). With the limited set of measurements and short DEER echo times presented here, it is difficult to define this interaction, but the appearance of an intermolecular peak suggests a significant surface expression of BtuB. Indeed, while BtuB is only natively expressed to a level of about 200 copies per cell, the pAG1 plasmid used here is capable of producing levels of BtuB equivalent to 40-50% of the level of the native porins (26). Experiments are currently in progress to better define this phenomenon.

An important finding reported here is the ability to perform double labeling and obtain DEER data from the periplasmic face of BtuB. Previous attempts to label the periplasmic surface of BtuB yielded no detectable signal (22), either due to a reduction of the nitroxide spin in the periplasm or the removal of the R1 side chain by the Dsb system. In the approach described here, the redox environment in the periplasm permits the maintenance of a spin label, and this may be due to both the use of cells at an earlier stage of log phase growth and the use of a *dsb* null mutant cell line.

An intriguing observation made here is that the behavior of BtuB on both the extracellular and periplasmic surfaces appears to be different in the native cell environment. On the extracellular surface, the addition of cobalamin to the V90R1-T188R1 pair promoted an interconversion between sub-peaks in the short-distance component, with a shift from 3.2 nm to a shorter distance at 2.4 nm (Fig. 4a). This shift is consistent with an upward motion of the core loop containing residue V90 towards residue T188 on barrel loop 2, and it is consistent with the unfolding of a short helical segment near this loop that is observed in the crystal structures (20). Remarkably, this shift is not observed in OM preparations, suggesting that cell environment is necessary for this substrate-dependent conformational change. The movement of this core loop is predicted by MD simulations of BtuB, but only in the presence of LPS (7). We are currently carrying out work to determine what features of intact cell system facilitate this conformational change.

On the periplasmic side, the Ton-box was examined in cells by measuring the distance between site D6 in the Ton-box and site Q510 on a periplasmic barrel turn. Compared to the reconstituted system, where the Ton-box was initially folded into the barrel and extended only with substrate addition (9), the Ton-box appears to be constitutively extended in the cell (Fig. 4b). Based upon MD simulations, the energy to unfold the Ton-box has been postulated to be dependent on the state of an ion-pair between site D316 in the barrel and R14 upstream of the Ton-box, which in turn depends on LPS interactions with the extracellular loops (7). Indeed, earlier EPR work using membrane reconstituted BtuB demonstrated that alanine mutations that break the D316-R14 ionic lock result in an unfolding of the Ton-box (27). In the MD study, the LPS was seen to induce a shift in the orientation of residue 14, which reduced the energy required to unfold the Ton-box. Whether this ionic interaction can account for the constitutively unfolded Ton-box remains to be tested. Nonetheless, these preliminary experiments indicate that a native behavior of BtuB requires the cellular environment.

## Conclusion

The approach described here demonstrates that pairs of cysteine mutants can be successfully maintained in a reduced state for spin labeling and pulse-EPR on both extracellular and periplasmic surfaces using mutants of the Dsb system. This will facilitate future work using EPR spectroscopy on BtuB and other OM proteins in the native cell environment. Our preliminary results obtained on both sides of the protein surface indicate that BtuB behaves differently in the cell than it does in a reconstituted system and may even be different than in an isolated OM preparation where transport does not occur. We anticipate that the ability to make these measurements on BtuB will enable measurements to unravel conformational rearrangements in the lumen of TBDTs and to elucidate the mechanisms of transport.

## Author Contributions

RKN provided the Dsb mutant cell lines. DAN and TDN performed all experiments, DAN carried out the data analysis, and DAN, TDN, RKN and DSC designed the experiments and wrote the manuscript. TDN and DAN contributed equally to this work.

## ACKNOWLEDGMENTS

We would like to thank Dr. Sarah B. Nyenhuis who provided assistance with some of the EPR measurements. This work was supported by a grant from the NIH (NIGMS, GM035215) to DSC. DAN was supported in part by a training grant from the NIH, T32GM080186.

## SUPPLEMENTARY MATERIAL

Supplementary material is available for this manuscript

